# *Listeria monocytogenes* Infection Alters Extracellular Vesicles Produced by Trophoblast Stem Cells to Confer a Pro-Inflammatory State

**DOI:** 10.1101/2022.04.19.488859

**Authors:** Jonathan Kaletka, Kun Ho Lee, Masamitsu Kanada, Jonathan W. Hardy

## Abstract

Placental immunity is critical for fetal health during pregnancy, as invading pathogens can be transmitted from the maternal blood to the fetus through this organ. However, inflammatory responses in the placenta can adversely affect both the fetus and the pregnant mother, and the balance between protective placental immune response and detrimental inflammation is poorly understood. Extracellular vesicles (EVs) are membrane-enclosed vesicles that play a critical role in placental immunity. EVs produced by placental trophoblasts mediate immune tolerance to the fetus and to the placenta itself, but these EVs can also activate detrimental inflammatory responses. The regulation of these effects is not well-characterized, and the role of trophoblast EVs (tEVs) in the response to infection has yet to be defined. The Gram-positive bacterial pathogen *Listeria monocytogenes* (*Lm*) infects the placenta, serving as a model to study tEV function in this context. We investigated the effect of *Lm* infection on the production and function of tEVs, using a trophoblast stem cell (TSCs) model. We found that tEVs from infected TSCs were immunostimulatory, activating macrophages to a pro-inflammatory state. Surprisingly, this activation made RAW 264.7 macrophages more susceptible to subsequent *Lm* infection. Increased susceptibility to infection has not previously been reported as an effect of EVs. Proteomic analysis and RNA sequencing revealed that tEVs from infected TSCs had altered cargo compared to those from uninfected TSCs. Together, these results suggest an immunomodulatory role for tEVs during prenatal infection.

## Introduction

The placenta is a remarkable organ in which the immune system plays a precarious role, balancing protective responses and potentially deleterious inflammation. The maternal decidua (the uterine tissue contacting the placenta) is an altered immune environment that permits the development of the semi-allogenic placental tissues and fetus; some pathogens exploit this immunosuppressed site, invading and replicating inside the placenta. Placental pathogens include viruses, parasites, and bacteria (1). One such organism is the Gram-positive bacterium *Listeria monocytogenes* (*Lm*) (2). This facultative intracellular parasite is the causative agent of listeriosis, an illness that affects approximately 1600 people annually in the United States, resulting in around 300 deaths (3). Listeriosis typically afflicts the immunocompromised, with pregnant people being especially at risk (4). Prenatal listeriosis can lead to spontaneous abortions, stillbirths, and birth defects, while pregnant mothers may show only mild symptoms (5). *Lm* is initially ingested with contaminated food such as deli meats, soft cheeses, and other dairy products. It escapes from the gastrointestinal tract and into the bloodstream, where it disseminates throughout the body and invades the liver, spleen, and the placenta (6). This pathogen has a well-characterized intracellular lifecycle which allows it to spread throughout host tissues, within monocytes and other cells. *Lm* enters the cell either by phagocytosis or by means of internalins, virulence factors that bind host surface proteins and induce uptake (7). Once in the cell, *Lm* is first contained in a phagosomal vacuole, which it lyses by means of the cholesterol-dependent cytolysin listeriolysin O (LLO), gaining access to the cytosol (8). In the cytoplasm, *Lm* scavenges the host for nutrients and replicates. Eventually, the bacterium hijacks and polymerizes host actin to create actin rockets, which facilitate intracellular motility and entry into neighboring cells, where it restarts this process (9, 10). Importantly, the ability of *Lm* to replicate intracellularly allows it to undergo cell-to-cell spread in trophoblasts, breaching placental barrier while minimizing exposure to the extracellular environment (6).

A coordinated cell-mediated immune response is critical to resolve *Lm* infection (11). A recently discovered mechanism that could play a role in mediating cellular immunity is intercellular communication mediated by extracellular vesicles (EVs). EVs are small (50-1000 nm) membrane-enclosed vesicles that are excreted by nearly every type of cell in the human body and across all domains of life (12). Two of the major types of EVs in eukaryotes have been historically designated as exosomes and microvesicles, which are differentiated based on how they are formed. Exosomes are smaller (50-150 nm) and are formed by the inward folding of the plasma membrane. Multiple vesicles are gathered and transported in multivesicular endosomes (MVEs), which fuse to the cell membrane and release the exosomes to the extracellular environment. Microvesicles, which have a wider range in size (100-1000 nm), are formed by the outward budding of the plasma membrane directly into the extracellular space (13).

EVs play a critical role in placental development and immune regulation during pregnancy. The number of EVs per volume of blood in a healthy mother greatly increases during pregnancy, and the majority of these EVs originate from fetal trophoblasts (14). Trophoblast EVs (tEVs) have been found to carry immunoregulatory molecules, presumably suppressing the immune response to allow for the successful development of the fetus (15). However, tEVs have also been experimentally shown to play a detrimental role during placental disease, such as preeclampsia (16, 17).

The role that EV-mediated communication plays during intracellular bacterial infections has only recently been explored with a select number of pathogens, and the role of tEVs during prenatal infection is unknown. Previous studies of *Mycobacterium*-infected macrophages showed that host EVs carry bacterial components such as RNA, proteins, and glycopeptidolipids (18–20). Other studies of *Salmonella enterica* serovar Typhimurium infections found similar results, in which EVs from infected macrophages carried *Salmonella* proteins and induced the production of pro-inflammatory cytokines through Toll-like receptor 2 (TLR2) and TLR4 dependent mechanisms (21). Additionally, mice treated with EVs from *S. enterica* infected cells generated antibodies against proteins found on the EVs, specifically *Salmonella* outer membrane proteins (22). EVs from *Lm-*infected macrophages carry bacterial DNA, and that this response is dependent on the DNA sensing cGAS-STING system (23). These findings show that EVs play a potential role in immune responses to intracellular bacterial infection. The purpose of the work presented here is to begin to decipher the role of tEVs produced in response to *Lm* placental infection.

We used an *Lm-*infected trophoblast stem cell (TSCs) system to model placental infections and tEV production and function. tEVs isolated from infected TSCs stimulated pro-inflammatory responses in recipient macrophage-like cells. Unexpectedly, we observed that RAW 264.7 cells became more susceptible to *Lm* infection after the tEV treatment. Using an untargeted proteomics approach, we found that tEVs from infected and uninfected TSCs had distinct protein profiles, with the infected tEVs containing more unique protein signatures than tEVs from uninfected TSCs. Ribosomal and other RNA binding proteins were increased in the tEVs by infection. However, in contrast to previous studies using macrophages, no bacterial proteins were found. RNA sequencing on the EVs revealed many mRNAs that were overrepresented in the tEVs from infected cells, including genes involved vasculogenesis and morphogenesis, processes involved in placental development. These data suggest that a unique mechanism is responsible for tEV-mediated immune modulation.

## Material and methods

### Bacterial cultures

*Listeria monocytogenes* 10403S bioluminescent strain 2C (Xen32) was used throughout the study (24). This strain has a *lux-kan* insertion in the *flaA* locus and has a four-fold increase in intravenous 50% lethal dose compared to wild type 10403S. It was grown in brain heart infusion medium (BHI) to mid-logarithmic phase for infection.

### Cell culture

Trophoblasts stem cells (TSCs) were originally isolated from C57BL/6 mice (25). They were grown in RPMI 1640 medium with GlutaMAX, 20% fetal bovine serum (FBS), and 1 μM sodium pyruvate, as well as 35 μg/mL fibroblast growth factor 4 (FGF-4), 10 ng/mL activin, and 1 μg/mL heparin to maintain TSC replication (26). RAW 264.7 and J774 cells were obtained from ATCC and were grown in RPMI medium with GlutaMAX, 10% fetal bovine serum (FBS), and 1 μM sodium pyruvate.

### Isolation of extracellular vesicles

107 TSCs in a 150 cm^2^ flask were infected with *Lm* at a multiplicity of infection (MOI) of 100, or treated with an equivalent volume of BHI. After 1 hour, the medium was replaced with medium depleted of EVs by centrifugation and containing 5 μg/mL gentamicin to ensure that there were no extracellular bacteria (27). At 24 hours of infection, the conditioned medium from the infected and uninfected TSCs was collected and centrifuged at 4000 x g for 20 minutes in 50 mL conical tubes. The supernatants were transferred to fresh conical tubes and centrifuged again at 4000 x g for 30 minutes. The supernatants were then filtered with a 0.22 μm filter using the Steriflip system. To collect large vesicles (L-tEVs), the filter was washed once with phosphate buffered saline (PBS), then 1 mL of PBS was repeatedly added to the top of the filter, which resuspended the tEVs from the filter. This preparation was then stored at -80° C. To collect the small tEVs (S-tEVs), the flow through from the filter was ultracentrifuged at 100,000 g for 2 hours. The supernatant was carefully removed so that there was about 0.5 mL left at the bottom of the tube, then 25 mL of PBS was added, and the preparation was ultracentrifuged again at 100,000 g for 2 hours. Once again, the supernatant was carefully removed, and the pellet was resuspended in an additional 1 mL PBS. The preparation was stored at -80° C.

### Transmission electron microscopy

108 tEVs were fixed in 2% paraformaldehyde for 5 min. 5 µL of the sample solution was placed on carbon-coated EM grids and tEVs were immobilized for 1 min. The grids were washed by transferring to five 100 µL drops of distilled water and letting it for 2 min on each drop. The samples were stained with 1% uranyl acetate. Excess uranyl acetate was removed gently with filter paper and the grids were air dried. The grids were imaged with a JEOL 100CXII transmission electron microscope operating at 100 kV. Images were captured on a Gatan Orius Digital Camera.

### Nanoparticle tracking analysis

tEV preparations were diluted 1:100 in PBS and were injected into a Zetaview machine (Particle Metrix). The Zetaview was set to a sensitivity of 89, a shutter speed of 300, and a frame rate of 30 frames per second. Cutoffs of 10 nm minimum and 1200 nm maximum were used.

### Listeria intracellular growth assay

J774 and RAW 264.7 macrophages were plated into a 24 well plate at 5 x 10^4^ cells/well. After 24 h, the cells were treated with 5 x 10^6^ tEVs from either uninfected or infected TSCs, or an equal amount of PBS. After another 24 h, the cells were washed three times with PBS. Medium with *Lm* was added at multiplicity of infection (MOI) of 10:1 colony forming units of *Lm* per cell. After 1 h, the wells were washed 3 times with PBS and medium with 5 μg/mL gentamicin was added. Bioluminescence images were taken at the given timepoints using an IVIS Lumina System (Perkin Elmer, Inc.), with 5 min of exposure and large binning, starting upon infection. The signal was quantified using Living Image software (Perkin Elmer).

### TNF-α quantification

J774 and RAW 264.7 macrophages were plated into a 24-well plate at 5 x 10^4^ cells/well. After 24 h, the cells were treated with 5 x 10^6^ of tEVs from either uninfected or infected TSCs, or an equal volume of PBS. After 24 h, the conditioned medium was collected and TNF-α was quantified by enzyme-linked immunosorbent (ELISA) assay from R&D Systems according to the instructions of the manufacturer.

### Fluorescence Microscopy

Flame-sterilized glass coverslips were placed into a 6-well dish. 10^5^ cells were seeded into each well. The cells were infected 24 h later with mid-log green fluorescence protein expressing *Lm* at an MOI of 100 as previously reported (28). After one hour, the media were replaced with medium containing 5 μg/mL gentamicin. At the listed periods post-infection, the cells were fixed with 4% paraformaldehyde and solubilized with 0.1% Triton X-100. The coverslips were treated with Rhodamine phalloidin (Invitrogen) for 30 min. The coverslips were then mounted to slides with DAPI Fluoromount-G (SouthernBiotech). The slides were imaged with Olympus Filter FV1000 confocal microscope and images were taken at 60x magnification.

### Proteomics

The protein profile of the purified tEVs was determined using untargeted mass spectrometry performed at the MSU Genomics Core Facility. Briefly, three independent EV preparations each of 10^9^ S-tEVs from uninfected and infected TSCs were lysed and the proteins were precipitated using acetone and digested with trypsin. Nanospray liquid chromatography with tandem mass spectrometry (LC-MS/MS) was used to determine the peptide profiles. The peptide data was analyzed using the Scaffold proteome software, which mapped the identified peptides back to the mouse and *Lm* references to determine the originating proteins.

### Gene Ontology (GO) Enrichment Analysis for proteomics

GO analysis was performed using the Gene Ontology Resource (http://geneontology.org/) and Protein Analysis Through Evolutionary Relationships (PANTHER) program to identify the biological processes of the proteins seen in the + Listeria S-tEVs (29). Additionally, protein interaction networks were generated using STRING program (30). Proteins that had twice the number of peptides identified in the + Listeria EV samples vs the – Listeria EV samples were used for the analysis.

### RNA sequencing

RNA extraction, RNA library preparations, sequencing reactions, and initial bioinformatics analysis were conducted at GENEWIZ, LLC. (South Plainfield, NJ, USA). Three independent EV preparations each of 10^9^ S-tEVs from uninfected and infected TSCs were used. Total RNA was extracted following the Trizol Reagent User Guide (Thermo Fisher Scientific). RNA was quantified using Qubit Fluorometer (Life Technologies, Carlsbad, CA, USA) and RNA integrity was checked with TapeStation (Agilent Technologies, Palo Alto, CA, USA). SMART-Seq v4 Ultra Low Input Kit for Sequencing was used for full-length cDNA synthesis and amplification (Clontech, Mountain View, CA), and Illumina Nextera XT library was used for sequencing library preparation. Briefly, cDNA was fragmented, and adaptor was added using Transposase, followed by limited-cycle PCR to enrich and add index to the cDNA fragments. The final library was assessed with Agilent TapeStation. The sequencing libraries were multiplexed and clustered on one lane of a flowcell. After clustering, the flowcell was loaded on the Illumina HiSeq instrument according to manufacturer’s instructions. The samples were sequenced using a 2x150 Paired End (PE) configuration. Image analysis and base calling were conducted by the HiSeq Control Software (HCS). Raw sequence data (.bcl files) generated from Illumina HiSeq was converted into fastq files and de-multiplexed using Illumina’s bcl2fastq 2.17 software. One mis-match was allowed for index sequence identification.

The raw PE reads sequencing data were uploaded to the Galaxy web platform, and we used the public server at usegalaxy.org to process the RNA-seq data (31). Briefly, PE reads were processed to trim sequencing adapter and low-quality bases using Trimmomatic (32). The clean PE RNA-seq reads were mapped to the mouse reference genome (Mus Musculus 10) using HISAT2 (33). Gene expression of mapped reads were then measured with featureCounts (34).

Differential gene expression analysis was performed using DESEq2 v1.32.0 in R version 4.1.1 (35). Genes with minimum 5 reads in at least 4 samples were filtered out, resulting in a total of 23,836 genes. Differentially expressed genes with p-adj < 0.05 were used to perform gene ontology analysis using the g:Profiler system (https://biit.cs.ut.ee/gprofiler/gost) (36). Biological processes with p-adj < 0.05 were considered significant. Volcano plot was generated using EnhancedVolcano package in R using fold change > 1 and p-value < 10-5 parameters (37).

### Mouse infections

All animal experiments were performed under IACUC-approved animal protocol 201800030 in accordance with BSL-2 guidelines established by Michigan State University Campus Animal Resources. Michigan State is an AAALAC International accredited institution. From 5 to 8-week-old BALB/c mice were obtained from Charles River Laboratories. They were housed in the Clinical Center Animal Wing at Michigan State University for two weeks to acclimate them. The mice were treated with either 200 μL of PBS, 10^8^ tEVs from uninfected TSCs, or 10^8^ tEVs from *Lm-*infected TSC through tail vein injection. After 24 h, the mice were infected with 10^4^ *Lm* through tail vein infection. At 96 h post infection, the mice were imaged using the IVIS imaging system as described previously (24, 38) and humanely sacrificed using cervical dislocation in accordance with approved procedures while the animals were anesthetized. The spleens were harvested, mashed, serial diluted, and plated onto BHI plates with 50 µg/mL kanamycin. Each spleen was diluted and plated in duplicate.

## Results

### Characterization of tEVs from infected TSCs

Trophoblast stem cells (TSCs) from C57BL/6 mice were used to model placental infections. TSCs were infected with *Lm* at a multiplicity of infection (MOI) of 100, and fluorescence microscopy was used to visualize the infection at 24 h post-infection (HPI), confirming that *Lm* can indeed infect these cells, as well as replicate and polymerize actin in this time frame (Fig. 1A). Additional infection of TSCs with a bioluminescent *Lm* strain confirmed that the bacteria are replicating at 24 HPI and continue to replicate beyond this time point (Supplemental Fig. 1A). These results show that TSCs are readily infected with *Lm,* albeit less efficiently than J774 macrophages or other professional phagocytes (27). At 24 HPI, the medium from uninfected and *Lm*-infected TSCs was collected and large tEVs (L-tEVs) were isolated by collecting the vesicles from the top of the 0.22 μm filter, while small tEVs (S-tEVs) were purified by ultracentrifugation at 100,000 x g. EV preparations have often been referred to as microvesicles and exosomes, but as these entities are formed by distinct processes and are not just differentiated based on their size, we will refer to our separated samples as L-tEVs and S-tEVs throughout this report (39). Transmission electron microscopy (TEM) on the tEV preparations show the distinctive round shape both vesicle preparations (Fig. 1B+C). The TEM images do show S-tEVs and L-tEVs that appear to be similar size to each other despite the different isolation methods. A possible explanation for this is that the L-tEV preparations have a more diverse population in size of EVs isolated. To determine if infection altered tEV production, we preformed nanoparticle tracking analysis on them using a Zetaview instrument (Supplemental Fig. 1B-E). We found that infection decreased the number of L-tEVs produced by TSCs but did not affect the number of S-tEVs (Fig. 1D). In addition, infection did not alter the size of either L-tEVs or S-tEVs (Fig 1E). Thus, *Lm* differentially affected tEV production, decreasing the number of L-tEVs isolated.

**Figure 1.**
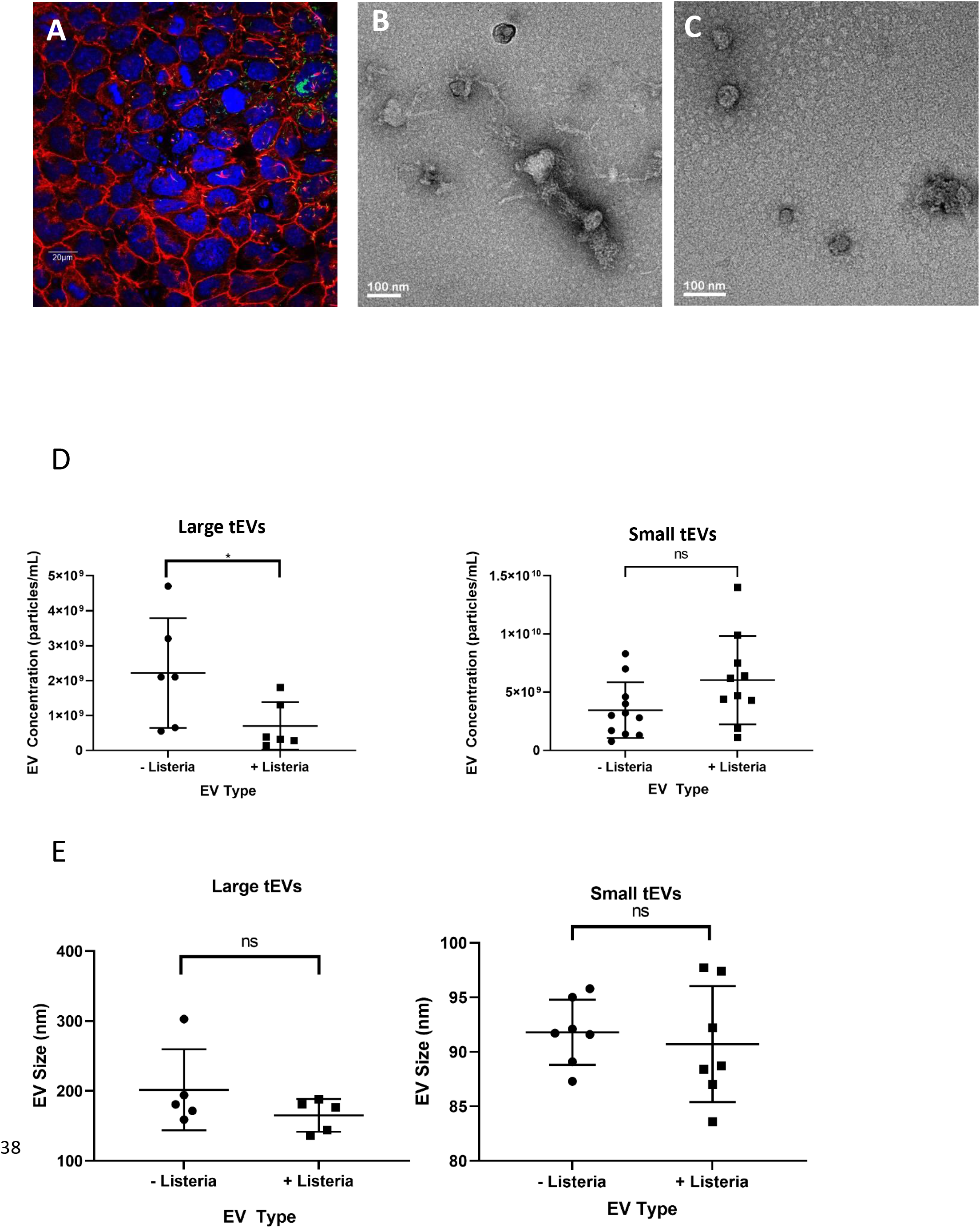
Extracellular vesicles from *Lm*-infected TSCs. (A) Trophoblast stem cells (TSCs) from C57BL/6 mice were infected with GFP-expressing *Lm* at a multiplicity of infection (MOI) of 100:1. At 24 hours post infection, the cells were fixed and stained with DAPI (Blue) and Rhodamine phalloidin (Red) which bind to DNA and polymerized actin, respectively. The cells were later imaged with an Olympus FluoView scanning confocal light microscope. The scale bar is 20 μm. (B, C) Transmission electron microscopy images of L- tEVs (B) and S-tEVs (C) from *Lm-*infected TSCs. (D, E) TSC derived EVs were analyzed by nanoparticle tracking analysis that gives the concentration and size distribution of the nanoparticles. The concentration (D) and mean size (E) of the tEVs with and without infection are given. Comparison using Student’s T-test, * *P*<0.05.

### tEV-mediated stimulation of macrophages

*Lm* infections lead to a pro-inflammatory response that is necessary to control the infection, with infected cells producing cytokines such as TNF-α (11). We hypothesized that tEVs can induce a similar pro-inflammatory response to *Lm* infection. To test this, we treated J774 and RAW 264.7 macrophage-like cells with 5 x 10^6^ tEVs derived from uninfected and infected TSCs, and after 24 h the TNF-α levels were measured using enzyme-linked immunosorbent assay (ELISA). We found that RAW 264.7 cells treated with L-tEVs or S-tEVs derived from *Lm-*infected cells resulted in the induction of TNF-α, while the treatment with PBS or tEVs from uninfected cells showed little TNF-α production (Fig. 2A). J774 cells, on the other hand, showed a significant increase of TNF- α only when treated with L-tEVs derived from infected cells, but not with S-tEVs from infected cells at the same concentration (Fig. 2B). Therefore, EVs isolated from *Lm*-infected TSCs have immunomodulatory effects in macrophages.

**Figure 2.**
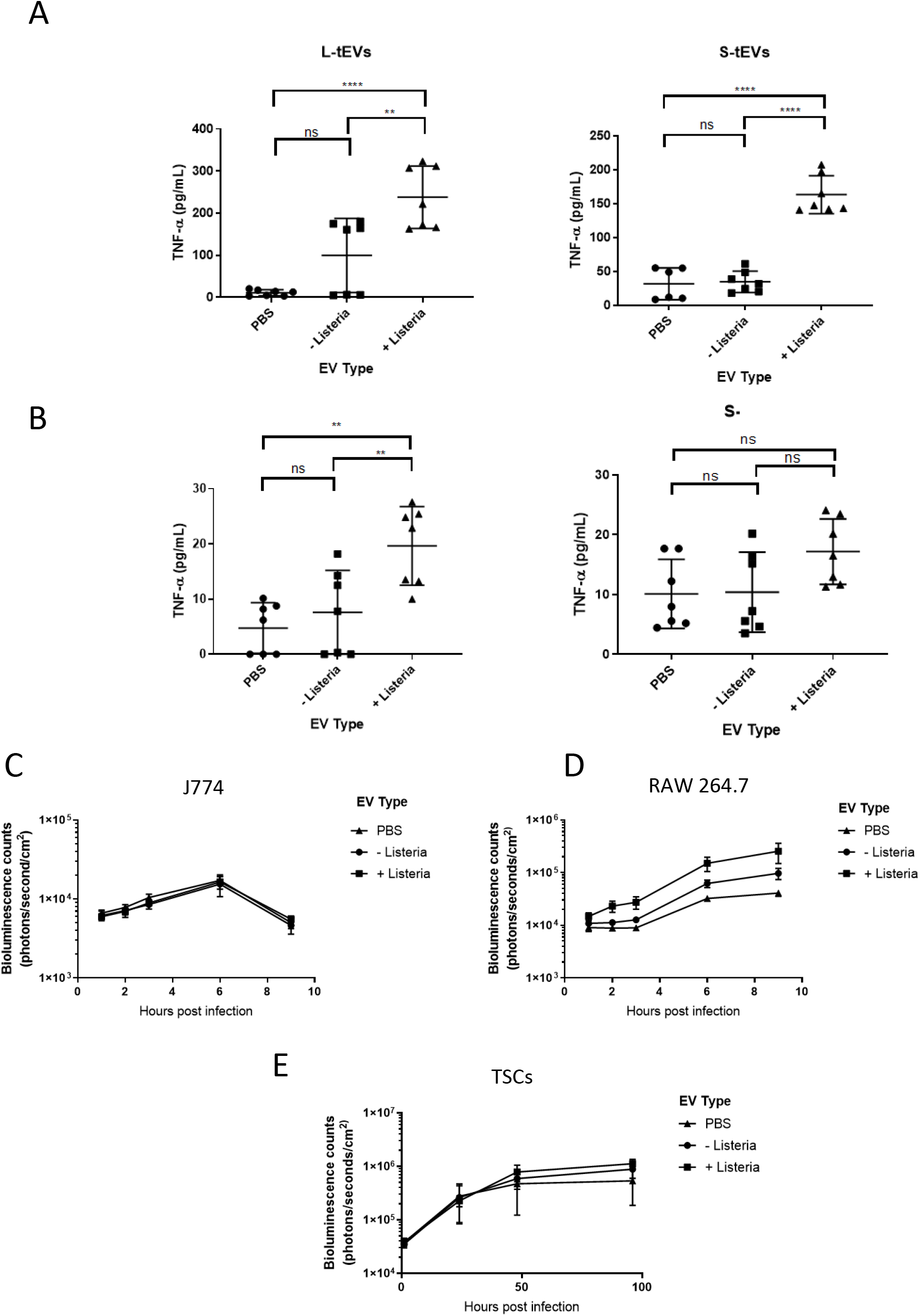
tEVs from *Lm-*infected TSCs activate RAW 264.7 cells. (A, B) 5x10^5^ RAW 264.7 (A) and J774 (B) macrophage-like cells were treated with 10^8^ tEVs from uninfected and *Lm-*infected TSCs and the production of tumor necrosis factor α (TNF-α) was measured using enzyme-linked immunosorbent assay (ELISA). (C) 5 x 10^5^ J774 cells, (D) RAW 264.7 cells, and (E) TSCs were treated with 10^8^ S-tEVs. After 24 h, the cells were infected at MOI=10 (C, D) or MOI=100 (E) with mid-log bioluminescent *Lm*. The cells were imaged using the PerkinElmer in vivo imaging system (IVIS). Comparisons for the TNF-α analysis was the Student’s T-test; ** *P<*.01, **** *P<*.0001.

EVs have been proposed as vaccines because of their ability to stimulate macrophages and induce antigen-specific memory (40–42). Additionally, macrophage activation causes increased resistance to *Lm* infection (43). We hypothesized that macrophages activated by the tEVs would become more resistant to *Lm* infection. We again treated RAW 264.7 and J774 cells with 5 x 10^6^ S-tEVs, and after 24 h infected with bioluminescent *Lm*. In J774 cells, no difference was seen in *Lm* growth (Fig. 2C). This result was expected since these tEVs failed to induce TNF-α production in J774 cells. However, we found that treatment with S-tEVs from uninfected TSCs increased the susceptibility of RAW 264.7 cells to infection. Surprisingly, treatment with tEVs from infected cells made the RAW 264.7 cells even more susceptible to infection (Fig. 2D). This result was unexpected, and as far as we are aware this is the first report of EVs inducing macrophages to become more susceptible to infection. Additionally, TSCs treated with tEVs also did not have a change in susceptibility to *Lm* (Fig. 2E), further suggesting that this is a cell type specific response to tEVs. Overall, we found that S-tEVs from infected TSCs activated RAW 264.7 cells and made them more susceptible to infection, and that this response was cell type specific, suggesting that tEVs are distinct from other EVs such as those from macrophages.

### Treatment of mice with tEVs

Our *in vitro* results suggest that tEVs from *Lm-*infected TSCs make macrophages more susceptible to *Lm* infection. However, there was also an increase in tEV-induced TNF-α production associated with infection of the source TSC cells. We therefore sought to determine the effect of tEV pre-treatment on *in vivo* infection, and whether tEVs would exacerbate the infection as indicated by the macrophage result or lessen the infection due to the induction of cytokines. We treated BALB/c mice intravenously (IV) with 10^8^ S-tEVs from uninfected or infected TSCs, or an equivalent amount of PBS. 24 h later, 10^4^ bioluminescent *Lm* were inoculated into the mice through IV injection (Supplemental Fig. 2A). At 72 hours post infection, the animals were imaged using IVIS imaging to quantify bacterial growth (Supplemental Fig. 2B). The spleens from the animals were also collected and serial plated to determine colony forming units. There was no statistically significant difference seen in the bioluminescence of the infection with any of the tEV conditions (Supplemental Fig. 2C), and although there was a slight decrease in spleen CFU recovered with tEVs from infected TSCs, this was also not statistically significant (Supplemental Fig. 2D). Taken together, tEV pretreatment does not appear to significantly affect the susceptibility of nonpregnant mice to *Lm* infection.

### Proteomic analysis of the effect on infection on tEVs

EVs carry a wide range of signaling molecules to deliver to recipient cells (44). We performed proteomic analysis to determine if *Lm* infection alters the protein profile of S-tEVs. Using shotgun tandem mass spectrometry, we found that there were many more unique proteins identified in S- tEVs from infected TSCs (331 proteins) compared to S-tEVs from uninfected cells (13 proteins). Additionally, in proteins that were shared between the two tEV groups (187 proteins), there were often more peptides identified in the infection condition, indicating higher amounts of that protein being transported in the tEVs from infected TSCs. Ribosomes, histones, and tubulin proteins are some of the categories where there were increased amounts in the infected tEVs. The full list of proteins that had at least a two-fold increase in peptide signature in the tEVs from the infected TSCs are listed in Supplemental Table 1. Meanwhile, only 4 proteins saw a two-fold increase in peptide signatures (Supplemental Table 2).

Gene ontology (GO) enrichment was performed to determine biological functions that were represented by the proteins that were increased in S-tEVs from *Lm*-infected TSCs. The main processes that were seen were related to ribosomes and translation, which is not unexpected given the many ribosomal and other RNA-binding proteins in these samples (Supplemental Table 2). A protein interaction map was created using the proteins that had at least a 2-fold increase in peptides in the S-tEVs from infected cells (Fig. 3). We found that these proteins have high levels of interaction with each other, and we can see clusters of ribosomal, cytoskeleton, and histone proteins. There were not enough proteins that were relatively increased in the tEVs from uninfected TSCs to perform GO analysis. Altogether, our results show that *Lm* infection does lead to different proteins loaded in tEVs, with more unique proteins seen in tEVs from infected cells. In contrast to other reports of EVs isolated from infected cells, no *Lm* proteins were detected in tEVs from any sample.

**Figure 3.**
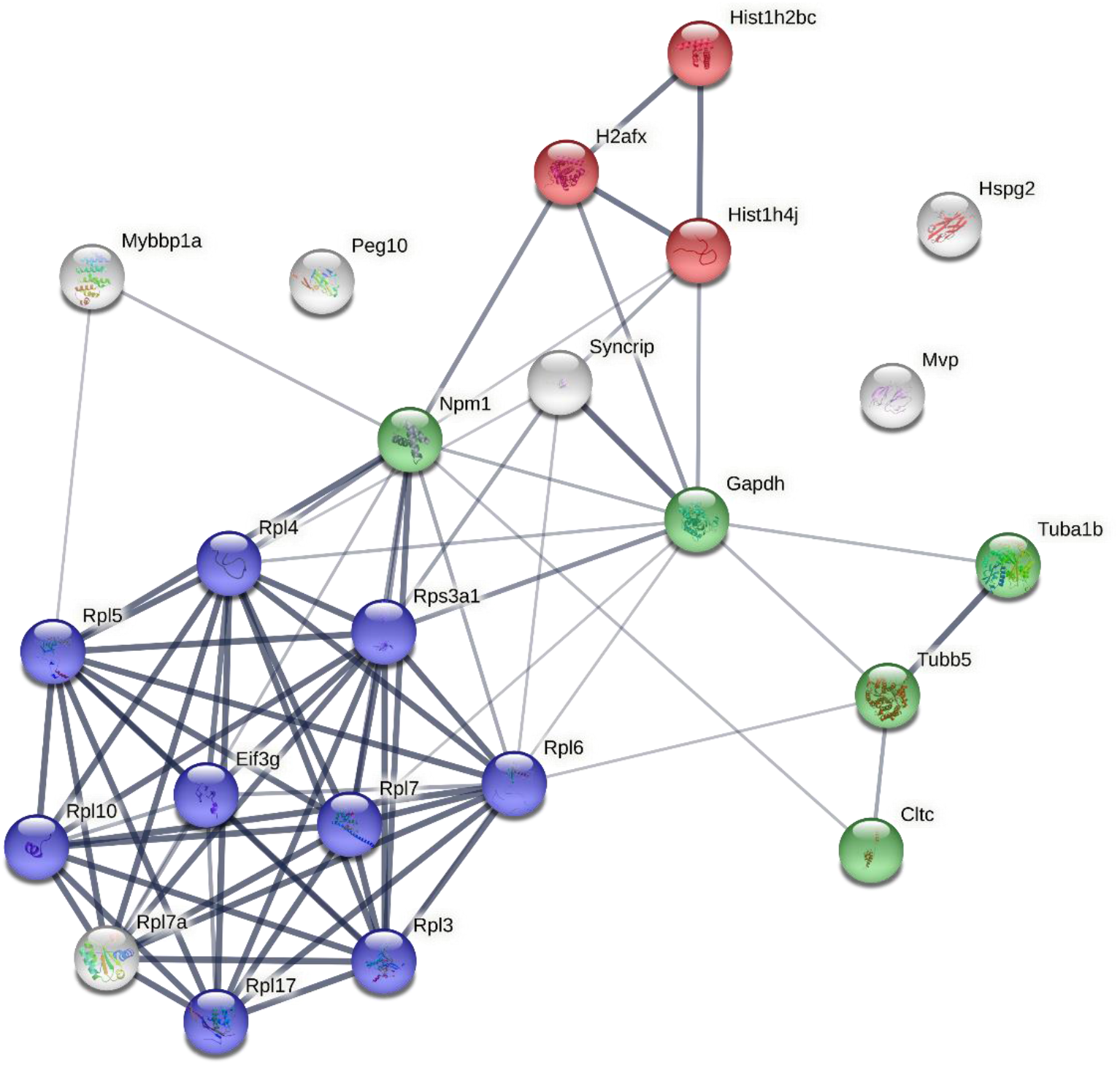
Protein interaction networks of proteins found in S-tEVs from *Lm*-infected TSCs. The proteins of the S-tEVs from uninfected and *Lm*-infected TSCs were determined by mass spectrometry. Protein interaction networks of the proteins identified in the S-tEVs from *Lm*-infected TSCs were generated using STRING program. The proteins involved in translation, microtubule cytoskeleton organization, and nucleosome core pathways are highlighted in blue, green, and red, respectively. Proteins that had twice the number of peptides identified in the *Lm*- infected tEV samples vs the uninfected tEV samples were used for the analysis. The thickness of the line represents the confidence of the interaction between the proteins.

### RNA sequencing on S-tEVs from Lm-infected TSCs

Our finding that S-tEVs from infected TSCs have increased amounts of RNA-binding proteins led us to believe that these EVs could also carry different RNAs. We performed RNA sequencing on S-tEVs from uninfected and *Lm-*infected TSCs (Fig. 4). We identified 22,836 genes in the mRNAs from the S-tEVs, with 68 genes being overexpressed in the S-tEVs from infected cells and 116 genes underexpressed in the S-tEV mRNAs from those cells (Fig. 4A). These differentially represented genes were used for GO analysis (Fig. 4B). Interestingly, two of the pathways upregulated in the genes of *Lm-*infected S-tEVs involved vascular development and morphogenesis, functions involved in placenta implantation and development. The pathways downregulated in the S-tEVs from infected TSCs involved metabolic processes. Overall, we found that *Lm* infection of TSCs altered the host mRNAs found in the S-tEVs. To our knowledge, this is the first reported findings of bacterial infections altering the host RNAs loaded into EVs.

**Figure 4.**
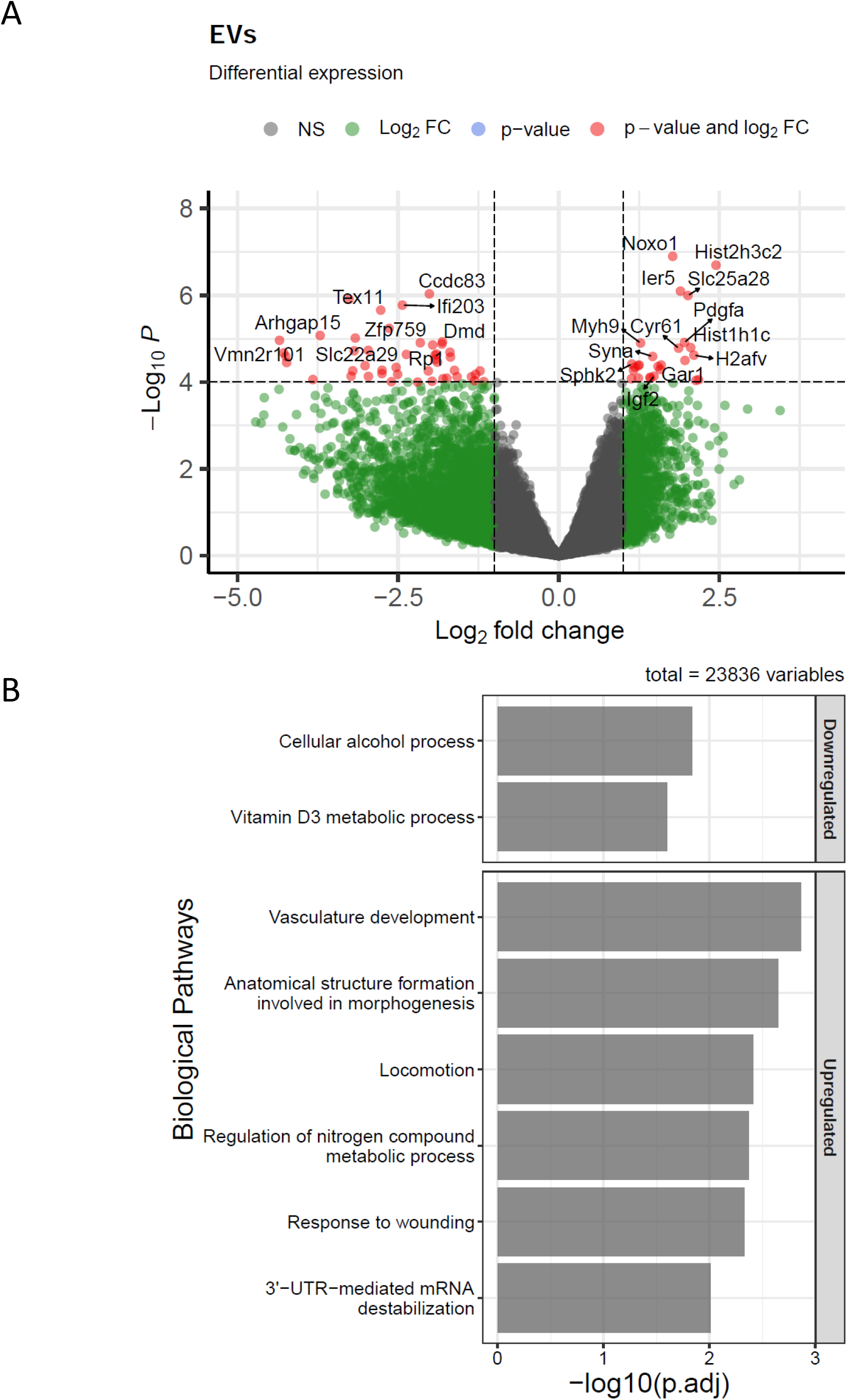
*Lm-*infected TSCs produce S-tEVs with altered RNA profiles. (A) Volcano plot of differentially expressed genes in S-tEVs from *Lm*-infected and uninfected TSCs. Red dots represent statistical significance (p-val < 10^-5^) and log2(fold change) greater or less than 1. Total variables represent the number of genes that were used to generate a volcano plot. (B) Gene ontology analysis of differentially expressed genes (p-adj <0.05) were used to investigate biological process pathways of downregulated and upregulated in *Lm-*infected cells using gProfileR.

## Discussion

EVs are the subject of exciting new research that offers the potential for novel approaches for diagnosis and treatment of many diseases. A primary function of EVs appears to be the stimulation of cell signaling pathways in recipient cells, including immune cells, activating them in some instances and dampening responses in others. This delicate balance can be altered during disease and infection. For example, certain cancers secrete EVs that suppress immune cell activity, allowing the malignancy to proliferate (45). Conversely, cells infected with intracellular pathogens secrete pro-inflammatory EVs that help to control infection, although EVs from infected cells may sometimes have the opposite effect (46).

During normal, healthy pregnancy, the number of EVs in the maternal bloodstream greatly increases, up to 60-fold compared to nonpregnant people (14). These EVs, produced by fetal trophoblasts of the placenta, have been shown to modulate immune responses but also can cause inflammation, leading to diseases such as the life-threatening preeclampsia. However, the function of placental EVs in prenatal bacterial infection remains unknown. Considering their large number and the above-mentioned effects on immune cells, placental EVs may play an important role in either ameliorating or exacerbating prenatal infection. We sought to study the effect of *Lm* infection on EV production and function by trophoblasts.

Modeling placental function *in vitro* is a challenge. Replicating trophoblast cell lines, such as BeWo and HTR8/SVneo, are convenient, but their genetic alterations conferring proliferation may compromise their response to infection (47). Primary human or mouse cells or explants are a superior representation of *in vivo* trophoblasts, but they are expensive and laborious to isolate and cannot be maintained in culture, requiring repeated isolation (48). TSCs offer an alternative between the two extremes. They are isolated from mouse blastocysts and are naturally replicating with the addition of growth factors, facilitating repeated controlled experiments while also maintaining many biological properties (25, 26).

As EVs are much smaller than eukaryotic cells or even bacteria, their isolation is complex and has been the subject of controversy. Currently, differential ultracentrifugation is the most commonly used method for isolating S-EVs. Low speed centrifugation removes cells and larger cell debris and subsequent high-speed ultracentrifugation (100,000 x g minimum) pools the tiny vesicles separate from the cells. Recent literature suggests that different isolation methods can affect the profile of EVs. Precipitation, density gradients, and filtration have all been used to isolate EVs, although each of these methods have their own positives and drawbacks (49). We chose to use differential centrifugation, paired with filtration to ensure preparations free of soluble protein and bacteria, because it is considered the gold standard method. It is important to note, though, that the EV isolation method can potentially influence the findings (39).

*Lm* is an intracellular pathogen that normally grows rapidly in macrophages *in vitro* unless the macrophage has been activated towards a pro-inflammatory M1 phenotype. For example, RAW 264.7 macrophage-like cells treated with IFN-γ are resistant to infection and kill intracellular *Lm* (43). Unexpectedly, we found that treatment with tEVs isolated from *Lm-*infected TSCs did not lead to increased *Lm* death; conversely, this treatment made the cells more permissible for growth of the bacteria. This result occurred despite the induction of TNF-α, which normally indicates stimulation of macrophages resulting in greater resistance to infection. As far as the authors are aware, this is the first account of an EV treatment that causes cells to become more susceptible to *Lm* infection. Importantly, this phenotype was only observed using RAW 264.7 cells, but not J774 cells. Both macrophage-like cell lines originate from BALB/c mice and have long been used in infection studies, although RAW 264.7 cells are from a male and J774 cells are from a female. J774 cells are more permissive for growth of *Lm*, and we expected activation by tEVs to reduce bacterial replication in these cells. The observation that these cells were equally permissive for *Lm* growth in all conditions was unexpected. The mechanisms of increased susceptibility in RAW 264.7 cells but not J774 cells remains of interest for future work. In addition, how the tEVs from infected TSCs stimulate recipient cells in the absence of bacterial products is under investigation and will require directed manipulation of the altered contents.

EVs represent a potential strategy by *Lm* to spread throughout the host by rendering recipient cells more susceptible to the bacteria. *Lm* invades humans through epithelial cells that line the intestines, and from there the pathogen relies on cell-to-cell spread to access the rest of the body. A previous report indicated that inhibiting EV formation attenuated *Lm* growth in the spleens of mice, further suggesting that *Lm* could be hijacking host EV function for its own benefit (23). That report found that EVs from *Lm-*infected cells carried *Listeria* DNA and activated the cGAS-STING pathway. This result is notable because this pathway leads to the release of type I interferons, which enhance *Lm* growth *in vivo* (50–52). Further studies will be needed to identify these pathways and define the role tEVs play during placental infection.

In mice, tEVs from infected TSCs detectably reduced infection, but the result was not statistically significant to p = 0.05. The reduction was modest, which could have been due to many factors. The number of tEVs administered (10^8^ tEVs), single vs. multiple tEV injections, whether or not the mice are pregnant, the timing and infectious dose of *Lm*, and the strain of mouse used may all be important, and we did not fully explore these parameters.

There were several interesting proteins detected in the tEVs. We were particularly interested to see an enrichment of ribosomal proteins in tEVs derived from infected cells. Other groups have also found ribosomal proteins when performing proteomic analysis of EVs (53–55). These studies usually focus on the RNA-binding aspects of these proteins as RNA has been heavily associated with EVs (13). However, the appearance of ribosomes in EVs could be independent of RNA. A potential explanation is that translation levels are increased during cell stress (such as an infection), which could lead to higher ribosome numbers, and eventually more ribosomal proteins in EVs. Another RNA-binding protein identified in our infection tEVs is PEG10. PEG10 is a Gag-like protein that is required for trophoblast differentiation and placental development (56). This protein can also selectively bind and load mRNA into exosomes, and these EVs alter the gene expression of the recipient cell (57). The ability of EV-associated PEG10 to alter gene expression could be an explanation for the altered behavior of cells treated with tEVs from *Lm*-infected cells, and this potential mechanism will be the focus of future studies. Surprisingly, these EVs lacked any *Lm* proteins in our analysis, contrasting previous EV studies during infection (19, 22, 58, 59). The majority of previous reports used macrophages as the initially infected cells, and the lack of *Lm* proteins may reflect reduced bactericidal mechanisms of trophoblasts compared to macrophages. This difference in protein processing and EV protein content could have important implications for the function of tEVs during pregnancy.

In addition to altered proteins found in the tEVs, we also saw a change in mRNAs during infection. One notable type of mRNAs observed in the S-tEVs from infected cells correspond to histone protein products. We also found histone proteins themselves in the tEVs from *Lm-*infected TSCs. Interestingly, histones have been identified in EVs in response to treatment with the gram-negative bacterial component LPS (53, 60). Additionally, histones have been identified in the bloodstreams of animal models of sepsis and patients with sepsis (61–63). These results in tandem with our findings suggest that packaging of histones, and potentially host cell DNA along with them, may be a mechanism to communicate infections. Further work is needed to determine if histones are responsible for the tEV activation of macrophages.

Some of the other genes that we saw overrepresented in the mRNAs from the infected S-tEVs involved vasculature development and morphogenesis. Vasculogenesis of fetal and maternal vessels are required steps for placenta implantation and development (64). Previous works with tEVs found that they recruit vascular smooth muscle cells and promote invasion of extravillious trophoblasts, key steps that are essential for remodeling the decidua surrounding the placenta (65, 66). EVs have also been found to play a direct role in implantation of the embryo (67), and inflammation is a key part of implantation (68). Additionally, morphogenesis of the placenta is required for the proper development of the organ (69, 70). One intriguing gene that had increased mRNA in the tEVs from infected is syncytin-A, which is responsible for the cellular fusion necessary for the development of the multinucleated synciotrophoblasts (71, 72). It is possible that *Lm* invasion of the placenta activates the release of tEVs required to carry out these processes, with the mRNAs housed in the vesicles acting on the recipient cells. Otherwise, the RNA profiles of the tEVs could represent the mRNAs being transcribed in the TSCs during infection, although the mRNAs we identified in the tEVs differ than those seen in human trophoblasts infected with *Lm* (73). Additionally, EVs have been previously found to contain mRNA profiles that differ significantly from the mRNAs in the cell of origin, and EV mRNAs can be translated in recipient cells (74, 75). More work is required to determine the exact function of tEV RNAs during infections.

Overall, we show that infection of TSCs with *Lm* alters tEV production and function in unexpected ways. tEVs from *Lm-*infected TSCs elicit immune activation in RAW 264.7 macrophage-like cells, and these cells are then less resistant to subsequent infection, which was unexpected. A multi-omics approach showed that *Lm* treatment greatly altered the components loaded into the tEVs, resulting in increased RNA and nucleic acid binding proteins and unique mRNAs in the tEVs from infected cells. The observation that there were no *Lm* proteins present in tEVs from infected TSCs suggests a host factor or factors altered in the tEVs is mediating the stimulation of target macrophages. The mechanism of this interaction is of great interest and is the subject of current investigation.

## Acknowledgements

We would like to thank the Institute for Quantitative Health Science and Engineering (IQ) at MSU for providing the facility and resources for executing this work. We gratefully acknowledge Dr. Melinda Frame and Dr. Alicia Withrow at the Center for Advanced Microscopy at MSU. Additionally, we also gratefully acknowledge Dr. Douglas Whitten at the Research Technology Support Facility Proteomics Core at MSU. Finally, we are most grateful for the team at Azenta/Genewiz, including Michael Bullard, Sabbir Siddiqui and Aleksandar Janjic, for their expert consultation and sequencing.

---

**Supplemental Figure 1.**
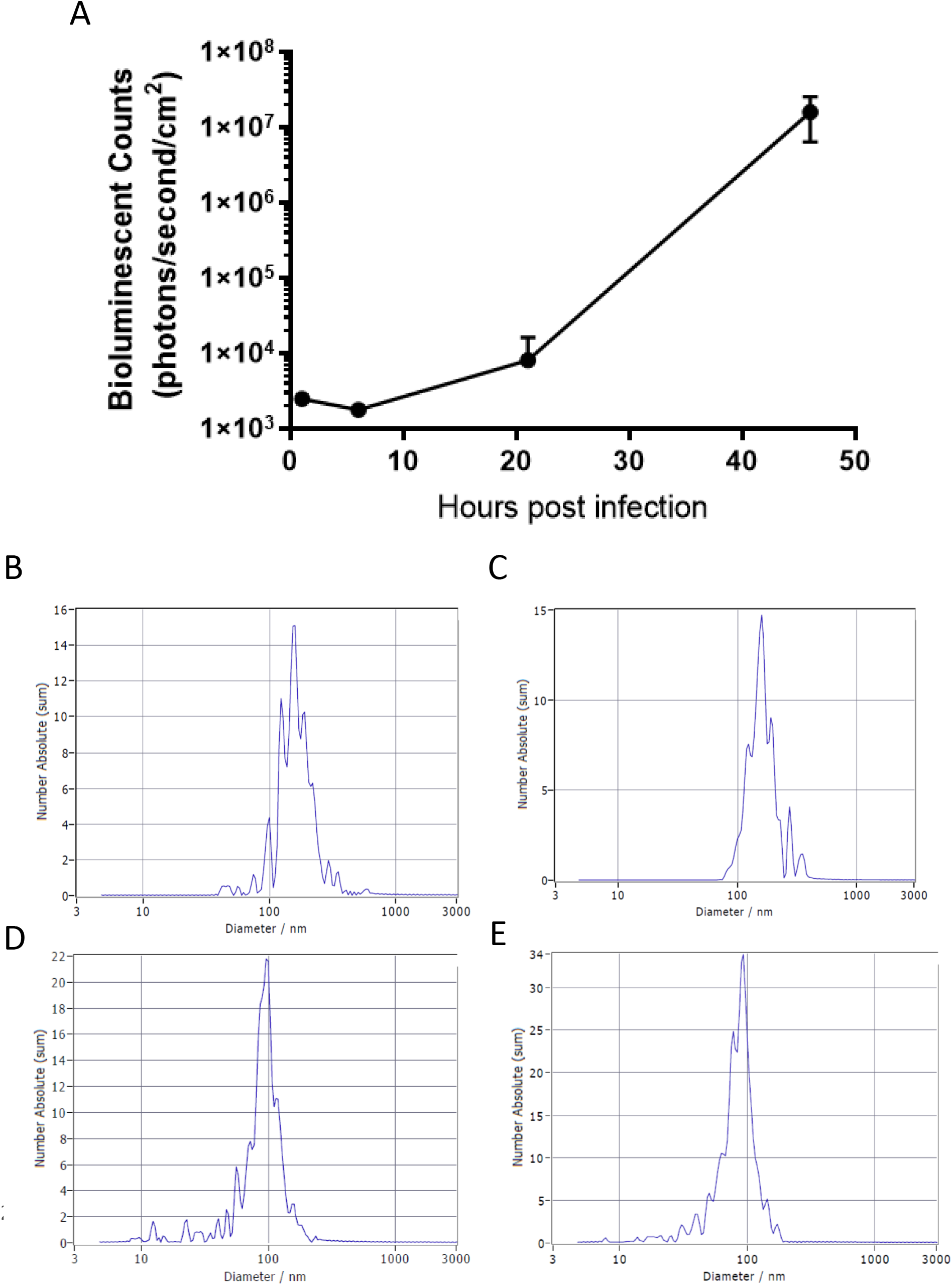
TSCs infected with *Lm* produce tEVs. (A) 5 x 10^5^ TSCs were infected at MOI=100 with mid-log bioluminescent *Lm*. The cells were imaged using the PerkinElmer in vivo imaging system (IVIS). (B-E) Example nanoparticle tracking analysis histograms giving the size distribution of tEVs. (B, C) Histograms for L-tEVs from uninfected (B) and *Lm-*infected (C) TSCs. Histograms for S-tEVs from uninfected (D) and *Lm-*infected (E) TSCs.

**Supplemental Figure 2.**
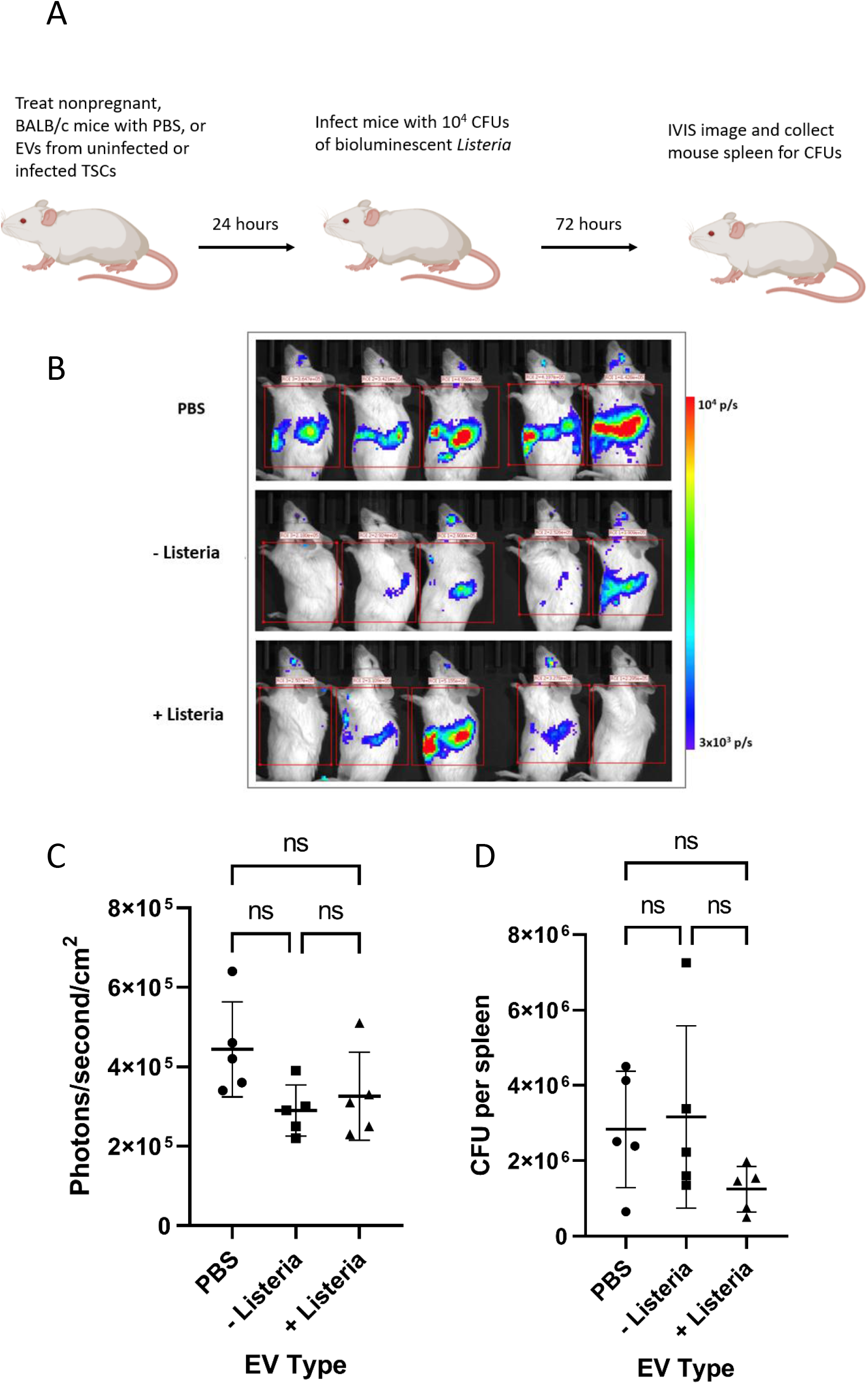
tEVs infection does not change mouse resistance to *Lm*. (A) Diagram of treatment of BALB/c mice with tEVs from TSCs before infection with bioluminescent *Lm*. Images made with BioRender. (B) IVIS images of mice 72 hours after infection. (C) Quantification of bioluminescence radiance signifying bacterial growth in the mice. (D) Bacterial growth measured by plate counting serial dilutions of spleens. Non-parametric one-way ANOVA (Kruskal-Wallis test) was used for comparison of the groups.

**Supplemental Table 1.** List of proteins that had twice the number of peptides identified in the S-tEVs from the infected TSCs vs. the S-tEVs from uninfected cells. Peptide counts were done in Scaffold Software.

| **Protein Name** | **- Listeria** | **+ Listeria** | **+ Listeria/- Listeria** |
| --- | --- | --- | --- |
| PEG10 | 0 | 6.33 | Infinite |
| 60S ribosomal protein L10 | 0 | 6.67 | Infinite |
| 60S ribosomal protein L3 | 0.33 | 11.33 | 34 |
| 60S ribosomal protein L5 | 0.33 | 9 | 27 |
| Myb-binding protein 1A | 0.33 | 7.67 | 23 |
| 40S ribosomal protein S3a | 0.33 | 5 | 15 |
| Nucleophosmin | 1 | 11 | 11 |
| Clathrin heavy chain 1 | 2.33 | 25.33 | 10.86 |
| 60S ribosomal protein L4 | 2 | 19.33 | 9.67 |
| Tubulin beta-5 chain | 2.33 | 22.33 | 9.57 |
| 60S ribosomal protein L6 | 1 | 8.67 | 8.67 |
| 60S ribosomal protein L7a | 1 | 8 | 8 |
| 60S ribosomal protein L17 | 1.33 | 10.33 | 7.75 |
| Major vault protein | 1 | 6.67 | 6.67 |
| Heterogenous nuclear<br>Ribonucleoprotein | 1.33 | 7.67 | 5.75 |
| Basement membrane-specific<br>heparan sulfate proteoglycan | 3.33 | 18.67 | 5.6 |
| 40S ribosomal protein 1A | 2 | 8.67 | 4.33 |
| Histone H2AX | 5.67 | 22.33 | 3.94 |
| 60S ribosomal protein L7 | 1.67 | 5.67 | 3.4 |
| Tubulin alpha-1B | 6 | 14.67 | 2.44 |
| GAPDH | 6 | 13.33 | 2.22 |
| Histone H2B | 11.67 | 23.33 | 2 |
| Histone H4 | 6.67 | 13.33 | 2 |

**Supplemental Table 2.**
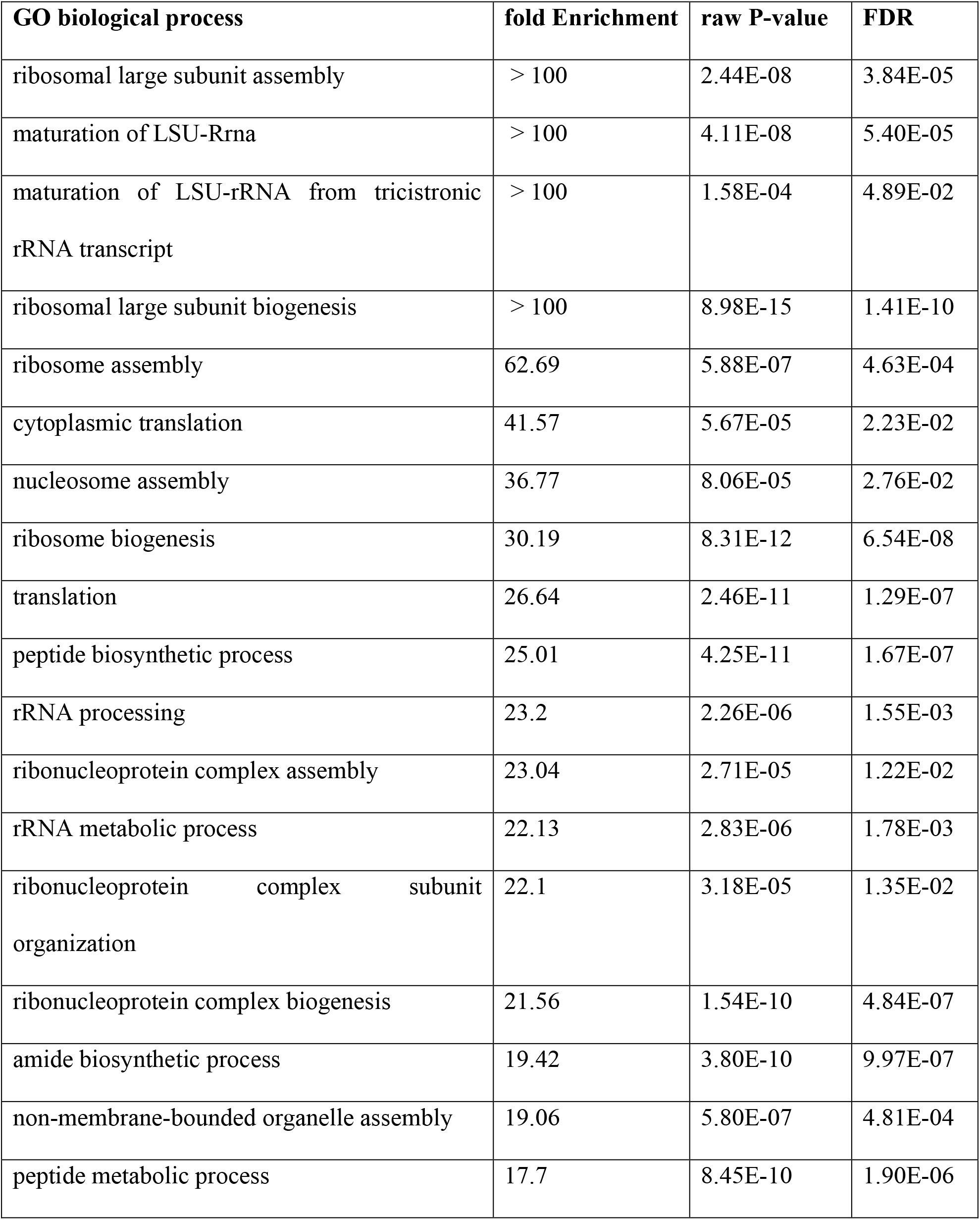
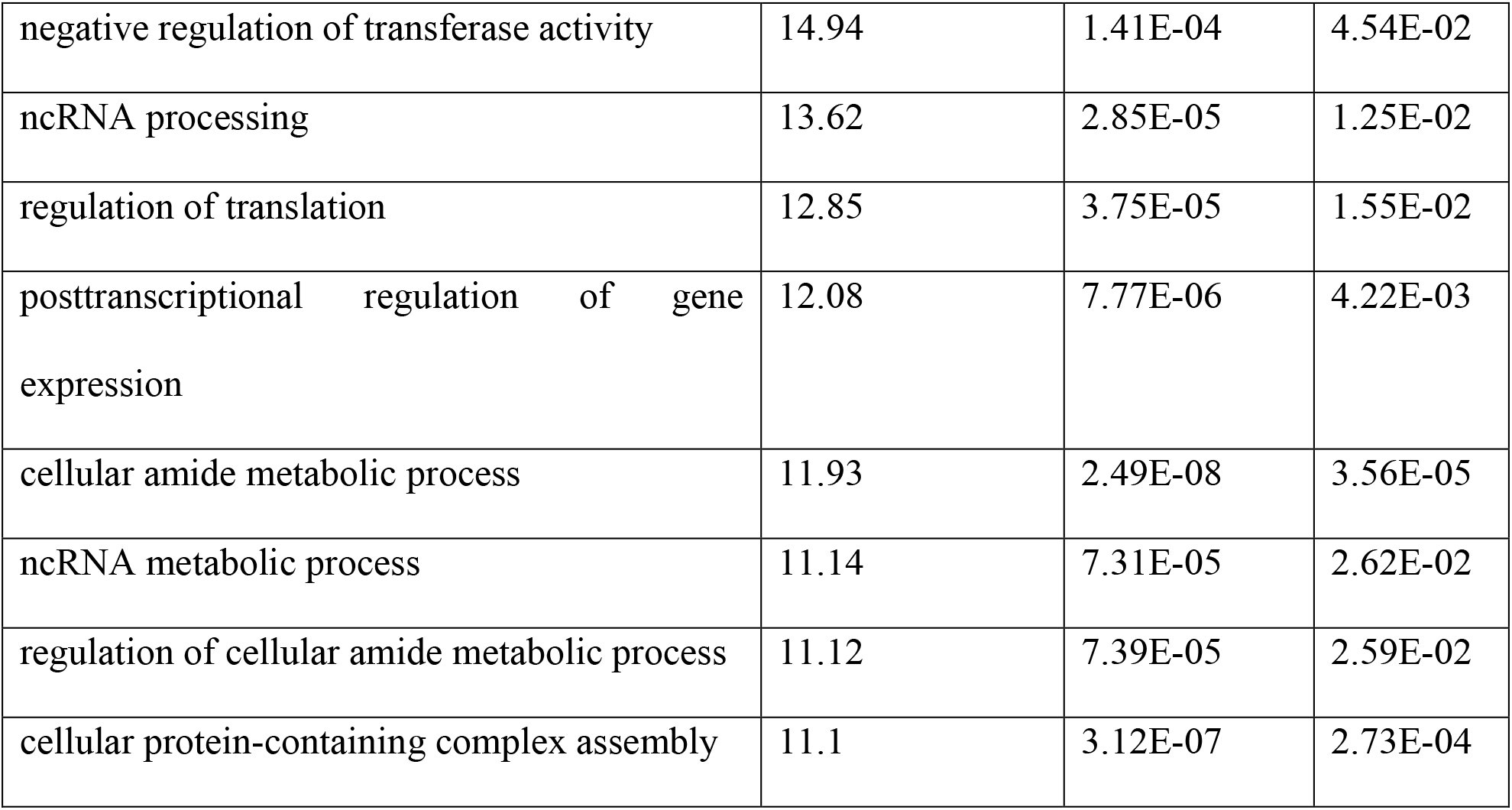
Gene ontology (GO) analysis with PANTHER was performed using the proteins identified in Table 1. The pathways that were associated with the proteins that were increased in tEVs from infected TSCs are listed, as well as the fold-enrichment scores, the P-values, and the false discovery rate (FDR).

| **GO biological process** | **fold Enrichment** | **raw P-value** | **FDR** |
| --- | --- | --- | --- |
| ribosomal large subunit assembly | > 100 | 2.44E-08 | 3.84E-05 |
| maturation of LSU-Rrna | > 100 | 4.11E-08 | 5.40E-05 |
| maturation of LSU-rRNA from tricistronic rRNA transcript | > 100 | 1.58E-04 | 4.89E-02 |
| ribosomal large subunit biogenesis | > 100 | 8.98E-15 | 1.41E-10 |
| ribosome assembly | 62.69 | 5.88E-07 | 4.63E-04 |
| cytoplasmic translation | 41.57 | 5.67E-05 | 2.23E-02 |
| nucleosome assembly | 36.77 | 8.06E-05 | 2.76E-02 |
| ribosome biogenesis | 30.19 | 8.31E-12 | 6.54E-08 |
| translation | 26.64 | 2.46E-11 | 1.29E-07 |
| peptide biosynthetic process | 25.01 | 4.25E-11 | 1.67E-07 |
| rRNA processing | 23.2 | 2.26E-06 | 1.55E-03 |
| ribonucleoprotein complex assembly | 23.04 | 2.71E-05 | 1.22E-02 |
| rRNA metabolic process | 22.13 | 2.83E-06 | 1.78E-03 |
| ribonucleoprotein complex subunit organization | 22.1 | 3.18E-05 | 1.35E-02 |
| ribonucleoprotein complex biogenesis | 21.56 | 1.54E-10 | 4.84E-07 |
| amide biosynthetic process | 19.42 | 3.80E-10 | 9.97E-07 |
| non-membrane-bounded organelle assembly | 19.06 | 5.80E-07 | 4.81E-04 |
| peptide metabolic process | 17.7 | 8.45E-10 | 1.90E-06 |
| negative regulation of transferase activity | 14.94 | 1.41E-04 | 4.54E-02 |
| ncRNA processing | 13.62 | 2.85E-05 | 1.25E-02 |
| regulation of translation | 12.85 | 3.75E-05 | 1.55E-02 |
| posttranscriptional regulation of gene expression | 12.08 | 7.77E-06 | 4.22E-03 |
| cellular amide metabolic process | 11.93 | 2.49E-08 | 3.56E-05 |
| ncRNA metabolic process | 11.14 | 7.31E-05 | 2.62E-02 |
| regulation of cellular amide metabolic process | 11.12 | 7.39E-05 | 2.59E-02 |
| cellular protein-containing complex assembly | 11.1 | 3.12E-07 | 2.73E-04 |

**Supplemental Table 3.** List of proteins that had twice the number of peptides identified in the S-tEVs from the uninfected TSCs vs. the S-tEVs from infected cells. Peptide counts were done in Scaffold Software.

| **Protein Name** | **- Listeria** | **+ Listeria** | **+ Listeria/-**<br>**Listeria** |
| --- | --- | --- | --- |
| Keratin type 1 cytoskeleton<br>(A6BLY7) | 11 | 3 | 0.27 |
| Trypsin | 66.67 | 23.33 | 0.35 |
| Lactotransferrin | 9.33 | 4 | 0.43 |
| Keratin type 1 cytoskeleton<br>(E9Q0F0) | 14.33 | 7 | 0.49 |

